# Exogenous dsRNA made accessible to Dicer by two eukaryotic RNA-dependent RNA polymerases

**DOI:** 10.1101/2025.04.26.650694

**Authors:** Marcello Pirritano, Johannes Buescher, Pauline Staubach, Thorsten Tacken, Yulia Yakovleva, Mark Sabura, Kristela Shehu, Sören Franzenburg, Marc Schneider, Martin Simon

## Abstract

Discrimination of self from non-self RNA is a critical requirement for any cell to respond to infections and to maintain cellular integrity. We report novel functions for two RNA-dependent RNA polymerases (RDRs) in *Paramecium* in the detection of exogenous RNA. In the RNA interference (RNAi) mechanism, RDRs are normally involved in the production of large amounts of secondary siRNAs in response to an initial primary siRNA. To characterize the function of RDRs in context of exogenous RNA recognition, we developed a novel dsRNA delivery system using dextran nanoparticles to deliver heteroduplex dsRNA to cells as food particles, mimicking the natural phagosomal entry pathway. Subsequent small RNA sequencing allows us to dissect siRNAs produced from exogenous RNA or RDR transcripts. Contrary to expectations of how dsRNA-induced silencing is triggered, our data show that Dicer is unable to directly cleave exogenous dsRNA while two RDRs are required for the initial steps of dsRNA-induced RNAi. Paradoxically, these two RDRs must replicate dsRNA to make it available for Dicer cleavage and primary siRNA accumulation. Our data also show that this system works efficiently also with exogenous ssRNA, although RDR2 is dispensable for ssRNA conversion. The function of RDRs in this protist is in contrast to that in animals, plants and fungi and extends the functional diversity of these polymerases. RDR-associated complexes appear to control the entry of food and symbiont-derived RNA into the RNAi machinery, enabling complex RNA interactions with the environment.

## Introduction

Any cell needs to have molecular mechanisms to recognize foreign RNA from pathogens while endogenous RNA needs to be tolerated. Double-stranded RNA (dsRNA) in particular was shown to trigger various mechanisms. In mammals, RigI-like receptors, PKRs and Toll-like receptors specifically recognize dsRNA and activate diverse immune responses [1, 2]. Nonvertebrates, in contrast, use RNA interference (RNAi) for antiviral defense, which involves the production of siRNAs to inactivate viral genomes and mRNAs. It has long been thought that mammals do not use antiviral RNAi, but a growing body of data supports the existence of RNAi in mammals; however, siRNA generation seems to be in competition with the interferon system. [3].

The decision whether dsRNA triggers RNAi or not is attributed to the activity of Dicer [4]. In nematodes, for example, exogenous dsRNA is rapidly cleaved by endogenous Dicer to functional siRNA [5]. Also in mammals, Dicer dissects the fate of dsRNA. Although canonical Dicer appears to be unable to cleave exogenous dsRNA, an oocyte-specific isoform lacking the helicase domain has increased activity on dsRNA [6]. Even more intriguingly, canonical Dicer in mammals has been implicated as a central mediator of antiviral responses by mediating a crosstalk to PKR and NF-kB [7].

Here, we analyze the cellular response to exogenous dsRNA in the ciliate *Paramecium*, in which RNAi can be easily induced by feeding dsRNA (reviewed in [8]). We provide evidence that this dsRNA is not the trigger molecule and is not processed directly by Dicer. Instead, exogenous dsRNA is first recognized by RNA-dependent RNA polymerases (RDR).

Eukaryotic RDRs are described for fungi, plants, animals, and also protists. In all these kingdoms except protists, the literature describes their general function in a process called ‘RNAi amplification’, which means the accumulation of large amounts of 2° siRNAs in response to a 1° siRNA that is cut by Dicer from an initial trigger molecule. This is demonstrated for yeast [9,10], plants [11], and also for *C*.*elegans* [12,13]. However, the nematode might not be representative for the animal kingdom, as RNAi in animals has been described to work without RDR amplification. Recent data suggest the existence of RDRs in animal genomes, but apparently with functions outside of siRNA amplification [14]. In contrast to detailed knowledge on RDRs in fungi, plant, and nematodes, functional analyses of protist RDRs are rare.

Questioning the role of exogenous trigger-mediated RNAi amplification in unicellular species, here we describe the function of two distinct RDRs in *Paramecium* in response to dsRNA. This particular method is an exciting mechanism to rapidly inactivate gene expression by simply feeding dsRNA-expressing *E*.*coli* [15] as described before in *C*.*elegans* [16]. Also similar to the nematode, 2° siRNAs have been demonstrated to occur, produced from the entire target mRNA [17]. While these appear to be Dicer independent, Dcr1 has been shown to be necessary for 1° siRNA production from dsRNA [18]: while the mechanistic role of RDR1 and RDR2 in 1° siRNA accumulation remains unclear [19].

## Materials and Methods

### Cell culture, synchronisation, phylogenetics, structural analysis

Wild-type and different mutant strains of *Paramecium tetraurelia* were maintained at 31 ° C in wheat grass powder (WGP) medium supplemented with *Klebsiella planticola* as a food source for culture maintenance or bacteria prepared for feeding experiments, and beta-sitosterol to promote growth as descriebd before [20]. Before experiments, cells were age-synchronized by daily isolation of single cells in fresh WGP medium and incubation at 31°C for 24 h, repeating isolation for a week. Cells were then incubated for two days in fresh medium to induce starvation and thereby autogamy. Autogamy was verified by DAPI staining.

For phylogenetic analysis, amino acids data were aligned with ClustalX and evolutionary history was inferred using the neighbor joining method with bootstraps relicates. Structural alignment of AlphaFold2 predicted structures [21] was carried out in ColAb v1.5.2 [22] and visualized using the Smith-Waterman 3D alignment method [23] in RSCB Protein Data Bank [24].

### Heteroduplex transcription, packaging and delivery

PCR products corresponding to the two RNA strands were created, harboring T7 promoters and terminators for subsequent *in vitro* transcription. *In vitro* transcription was carried out by the HiScribe T7 High Yield RNA Synthesis Kit from NEB following the manufacturer’s instructions. Prior to DNAse I treatment and phenol/chloroform extraction, the produced RNA strands were annealed to a heteroduplex dsRNA by heating the reaction to 85°C for 10min followed by slow cooling to 4°C over a period of 45min.

Annealed heteroduplex dsRNA was mixed in aqueous solution at 1:2 mass ratio with DEAE dextran 20 (TdB Labs, Uppsala) to facilitate nanoparticle formation using electrostatic interactions. The size of the particles and the net charge were measured using a Zetasizer Ultra (Malvern Panalyitcal) and the particles were visualized by transmission electron microscopy using a JEOL JEM-2100 in bright field mode with a slow-scan charge-coupled device camera at 200kV operating voltage. To facilitate heteroduplex nanoparticle uptake in *P. tetraurelia*, positively charged particles were adsorbed onto the negatively charged surface of *E. coli* by harvesting an bacterial overnight culture, washing the bacteria twice in 1xPBS and mixing 1ml of resuspended *E. coli* with 50 ul nanoparticle suspension and incubating the mixture for 30min at 37°C under constant shaking. Bacteria were then mixed with 20ml WGP medium. *Paramecia* were grown in this medium for 24h at 27°C before RNA extraction.

### RNA feeding using *E. coli*

For single strand RNA application, *E. coli* cells of the HT115 DE3 strain were transformed with a plasmid that harbors a fragment of the *ND169* gene under control of a T7 promoter and terminator. To prepare single-strand RNA feeding medium, a culture of the transformed *E. coli* was inoculated using an overnight culture and grown to an OD of 0.4 before IPTG was added to induce ssRNA production. ssRNA production was induced for 2.5 h until the bacteria were harvested by centrifugation and transferred to WGP medium. Paramecia were cultivated in the prepared feeding medium for 48h at 27°C and then harvested for RNA extraction. DsRNA feeding using the same *E. coli* strain was performed, using a L4440 vector containing a fragment of the *ND169* gene as described before [25].

### RNA extraction, sRNA-seq, and biochemistry

RNA was extracted from 50.000 cells using TRI Reagent (Sigma-Aldrich). RNA was size-selected using a UREA-MOPS PAGE and enriched for RNA molecules between 18-30nt. Size-selected RNA was used as an input for sRNA library preparation using the NEBNext Small RNA Library Prep Kit for Illumina from NEB according to the manufacturer’s instructions following the size selection by TBE-PAGE protocol. To distinguish sRNAs with different biochemical properties, mainly their 5’-phosphorylation status, size-selected sRNA was treated in different ways to create libraries specific for certain modifications. For 5’-mono-phosphate specific libraries, the NEBNext Small RNA Library Prep Kit for Illumina from NEB, based on classical ligation, was used. For 5’-mono- and tri-phosphate smallRNAs, size-selected RNA was treated with the CapClip enzyme (Cellscript) and then converted into a library as described, or untreated size-selected RNA was subjected to library preparation by using a template switch-based library kit (D-Plex Small RNA-seq Library Prep Kit for Illumina from Hologic Diagenode). For 5’-tri-phosphate specific libraries, size-selected RNA was first Terminator digested (Epicentre) to digest 5’-mono-phosphate RNA. After extraction with acid phenol, the remaining RNA was treated with the CapClip enzyme (Cellscript) before being subjected to the NEBNext Small RNA Library Prep Kit for Illumina from NEB. Successful treatment for biochemical discrimination of RNA was validated by using RNA spike-in oligos with known 5’-phosphorylation status, controlled using a UREA-MOPS PAGE. A DNA spike-in oligo was used as a loading control. Libraries were multiplexed and sequenced on the Illumina NextSeq.

### Data processing and analysis

Reads were de-multiplexed and quality- and adapter-trimming was carried out using the TrimGalore tool, which uses Cutadapt [26, 27]. Quality of reads was evaluated using the FastQC/MultiQC tool [28, 29]. Only reads between 18-30nt in length were maintained. For heteroduplex analysis, reads were mapped to the two heteroduplex sequences simultaneously, using the Bowtie mapper allowing no mismatches or multi-mapping [30]. Reads mapping on the individual strands of the heteroduplex in either sense or antisense orientation were quantified using in-build Tools of the Geneious Prime 2024.0 Software (https://www.geneious.com).

For analysis of untemplated nucleotides, a modified version of a custom snakemake pipeline was used (https://www.github.com/greenjune-ship-it/untemplated-nucleotides-search). In short, sRNA reads were mapped to the template sequence without mismatches using Bowtie [30]. Then, reads mapped to the sequences were extracted, sense and antisense directed reads were separated and sequence logos for each read length were calculated with the weblogo tool [31]. Reads extracted this way were considered not containing untemplated nucleotides. In a second iteration, reads not mapped were trimmed by removing a single base from the 3’-end using Cutadapt [27] and mapped to the template sequence again. Reads mapped in this iteration were processed the same way as described above and were considered to carry a single untemplated nucleotide. This process was repeated a total of four times, allowing for the analysis of up to three untemplated nucleotides. Overlap analysis of siRNAs was performed using the siRNA and piRNA overlap signature prediction tool ([32]). All sequencing data are available in the ArrayExpress database (http://www.ebi.ac.uk/arrayexpress) under accession number MTAB-14931.

### RDR localization

For GFP-assisted localization of RDR1 and RDR2, young *Paramecium* cells were transfected by macronuclear injection of a linearized plasmid containing the open reading frame of either RDR fused with an N-terminal GFP. To enhance visualization of RDR localization, GFP-positive cells were used to perform an immunofluorescence assay using custom anti-GFP antibodies.

## Results

### 1° and 2° siRNAs in *Paramecium* are 5′-monophosphorylated

We first evaluated the biochemical properties of siRNAs. This is necessary since RDR-dependent amplification mechanism are diverse: in plants, they produce long dsRNA and subsequent Dicer cleavage creates double stranded siRNAs carrying a 5′-mono-phosphate (5′-P) [33], whereas 2° siRNAs carrying a 5′-tri-phosphate (5′-PPP) in *C*.*elegans* as a result from *de novo* transcription by RDR. [34].

In *Paramecium*, 2° siRNAs presumably produced by RDR2, map to mRNA outside of the feeding region [17]. To clarify whether RDR associated siRNAs in *Paramecium* produce any other than 5′-P siRNAs, we treated RNA samples from RNAi against the *ND169* reporter gene either with Cap-Clip pyrophosphatase (converting any phosphorylation in ligatable 5′-P), or by Terminator digestion with subsequent Cap-Clip treatment (digest of 5′-P RNA with subsequent conversion of any remaining RNAs into 5′-P RNA). Both treatments were then subjected to classical, 5′-P specific ligation based library preparation. In addition, a 5′-phosphorylation independent template switch based library preparation was carried out. Fig.1A shows siRNA coverage of the *ND169* gene for all treated RNAs, highlighting the fragment used for dsRNA synthesis (corresponding to 1° siRNAs) with 2° siRNAs being found outside of the highlighted area. Comparing the different treatments to the 5′-P specific library, we cannot identify any differences: abundant 1° siRNAs in the dsRNA region and a low coverage of 2° siRNAs outside this regions in all samples (ratio 1°/2° approx. 2-5 percent (Suppl. Fig. 1). Please note the low read number in the Terminator-treated sample. This library was almost invisible during gel extraction, indicating that only remnants of RNA were ligated which is supported by the read length distribution Suppl. Fig. 1B). Therefore, we can rule out that we miss a high number of 2° siRNAs by 5′-P dependent ligation procedures and that 2° siRNAs are of much lower abundant than 1° in *Paramecium* (Suppl. Fig. 1C).

**Fig 1.**
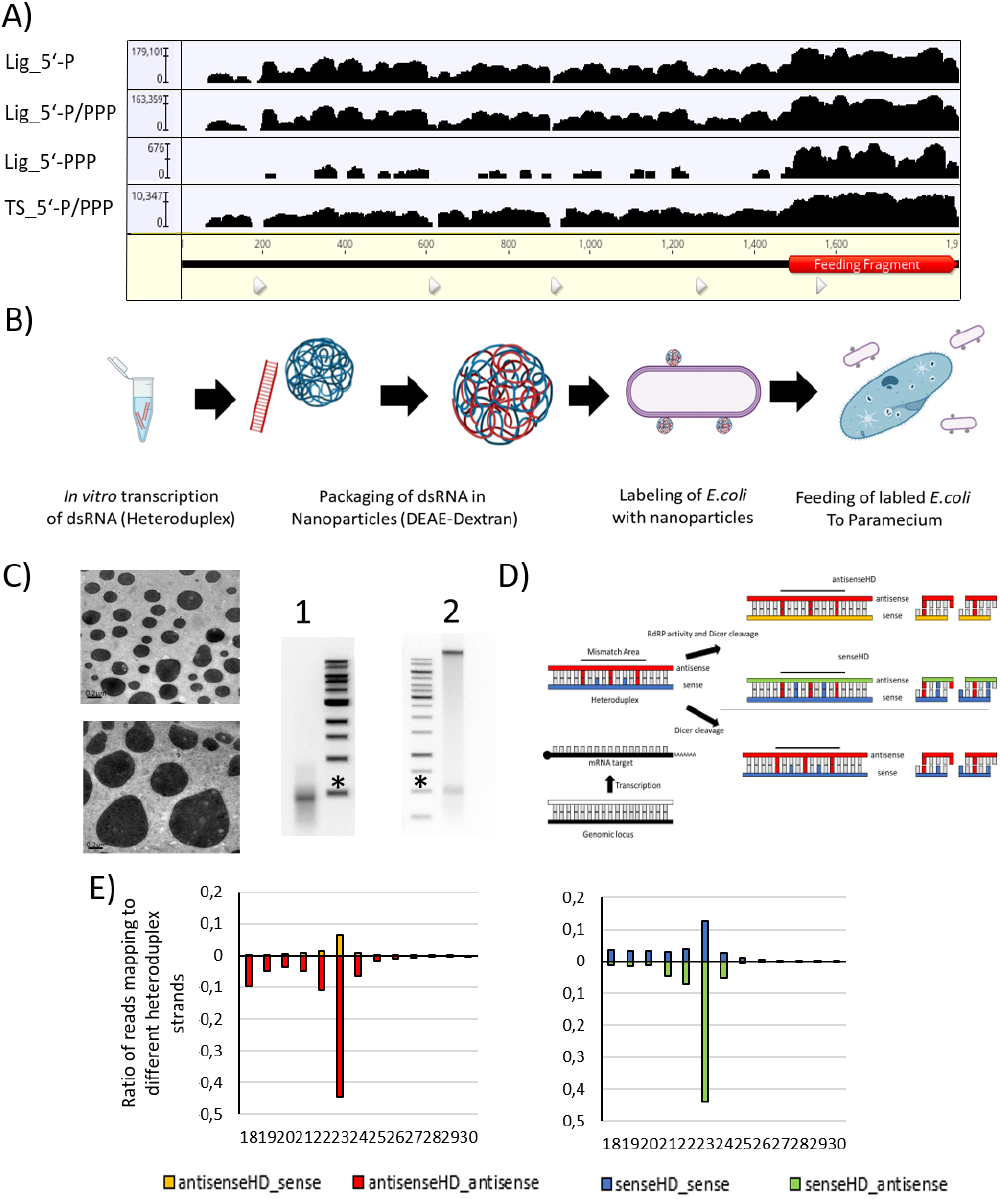
A) Coverage of 23nt siRNA reads produced from dsRNA feeding against the *ND169* gene. The feeding fragment is highlighted in red (1° siRNAs). siRNAs outside of the feeding fragment region are 2° siRNAs. Libraries enriching RNAs for individual 5’-phosphorylation status are: P - mono-phosphorylation; PPP - tri-phosphorylation. Lig - ligation-based library preparation, TS - template switch based library. Note the logarithmic coverage scale B) Workflow of heteroduplex application from *in vitro* transcription, nanoparticles assembly, labeling of *E. coli* and application of labeled bacteria to *P*.*tetraurelia*. C) Right: Transmission electron microscopic image of nanoparticles. Scale bars are 0.2µm. Left: Native Gel electrophoretic analysis annealed dsRNA before nanoparticle formation (lane 1) and after particle formation and adsorption to *E. coli* (lane 2). RNA in lane 2 was re-isolated from living bacteria. The high molecular band corresponds to the bacterial gDNA. Asterisks indicate 500bp Marker bands. D) Scheme of heteroduplex strands (blue and red) in association to the endogenous target. Mismatches between heteroduplex strands as well as the endogenous sequences are highlighted. RDR activity on the heteroduplex leads to new strands (orange, green). Lack of RDR activity leads to generation of siRNAs solely based on original heteroduplex strands. E) Mean read length distribution of sRNAs. Abundance of reads corresponding to sense and antisense strands of the original strands (blue and red, respectively) and to the RDR-dependent products (green and orange) are shown. Mean distribution of four different replicates is displayed.

### 1° siRNAs are a mixture of applied dsRNA and RDR products

In order to elucidate the role of RDR1 and RDR2, we decided to apply a specific heteroduplex dsRNA *in vivo*. We have chosen this method rather than analyzing *in vitro* enzymatic activity with purified enzymes, which often produce artifact activities. Since direct production of heteroduplex in *E*.*coli* is impossible due to rapid recombination of the two nearly identical parts in the plasmid, we annealed single-stranded *in vitro* transcripts and incorporated them into dextran nanoparticles (Fig. 1B,C). To guarantee cell entry through the natural phagosomal pathway, the dextran particles were adsorped to *E*.*coli* cell surface and then fed to paramecia (Fig. 1B). Successful re-isolation of intact dsRNA from *E*.*coli* demonstrated the stability of the trigger molecules (Fig.1C). After feeding the manipulated E.coli to paramecia, small RNAs were sequenced and mapped to the individual template strands of the heteroduplex sequence containing the mismatches. The heteroduplex dsRNA is designed with mismatches (i) between the two strands and (ii) to the target mRNA. This allows dissection of which strand an siRNA is derived from, one of the originally applied strands (blue/red), their RDR transcripts (green, yellow), or from a genomic RNA (black/white) (Fig. 1D and Suppl. Fig. 2A). Figure 1E shows that we can detect siRNAs not only from the original strands (blue and red) but also from RDR transcripts of both (green and yellow). The experiments were repeated four times with almost identical results (Suppl. Fig. 2B). We conclude that 1° siRNAs in *Paramecium* are a mixture of endogenous RDR products and exogenously applied RNA. RDR products are not in excess, which argues against massive amplification. We can also detect a very low number of antisense reads originating from mRNA, corresponding to 2° siRNAs (Suppl. Fig. 2B).

Length distributions of siRNAs show a 23nt peak with a preference for antisense siRNAs, regardless of whether they originate from the original dsRNA (red) or from RDR products (green) (Fig. 1E). To evaluate whether this antisense preference could be due to the introduced mismatches, we performed an experiment with a switched heteroduplex in which the additional mismatches are not in the antisense strand, but in the sense strand of the dsRNA. Suppl. Fig. 3 shows that antisense siRNAs are also more abundant than sense siRNAs in this RNA. We conclude that stabilization of antisense siRNAs occurs independently of the origin (exogenous or RDR product) of the strand.

### Sense siRNAs carry non-templated Uridines

To get deeper insight in the accumulation of antisense RNAs, we screened the sRNAs resulting from heteroduplex feeding for sequence characteristics. Suppl. Fig. 4 shows that we cannot identify a preference for nucleotides in any sRNA from the heteroduplex. This distinguishes feeding associated small RNAs from other sRNA produced by Dicer1 in *Paramecium* as e.g. transgene-induced siRNAs show a 5′-U preference [35]. The sequence logo analysis of normal dsRNA feeding also did not reveal any nucleotide preferences, so we can rule out putative influences of the mismatches in the analysis (Suppl. Fig 4 B). Classical non-heterduplex feeding analysis allows in addition for an analysis of overlapping reads. Suppl. Fig. 4B shows a peak for 21nt overlap probability which would fit to 23nt Dicer cuts with 2nt overlaps and this is consistent with previous reported that 1° siRNAs in *Paramecium* depend on Dcr1 [17, 18].

Our analysis of non-templated nucleotides indicates higher levels with sense siRNAs (Fig. 3A). An unbiased analysis of these non-templated nucleotides using the custom pipeline by sequence logo analysis identifies these non-templated nucleotides as uridines that extent 23nt siRNAs. Fig. 3B shows that these occur exclusively to sense siRNAs, regardless of their source (RDR product or exogenous strand). Since oligouridinylation of sRNAs is mostly associated with a targeting for degradation [36], we hypothesize that sense siRNAs are targeted for degradation, explaining the observed antisense bias, while antisense siRNAs are stabilized in Piwi. Our data do not implicate any difference between the two types of antisense and sense strands, respectively, and suggest that both duplexes are processed in the very same manner.

**Fig 2.**
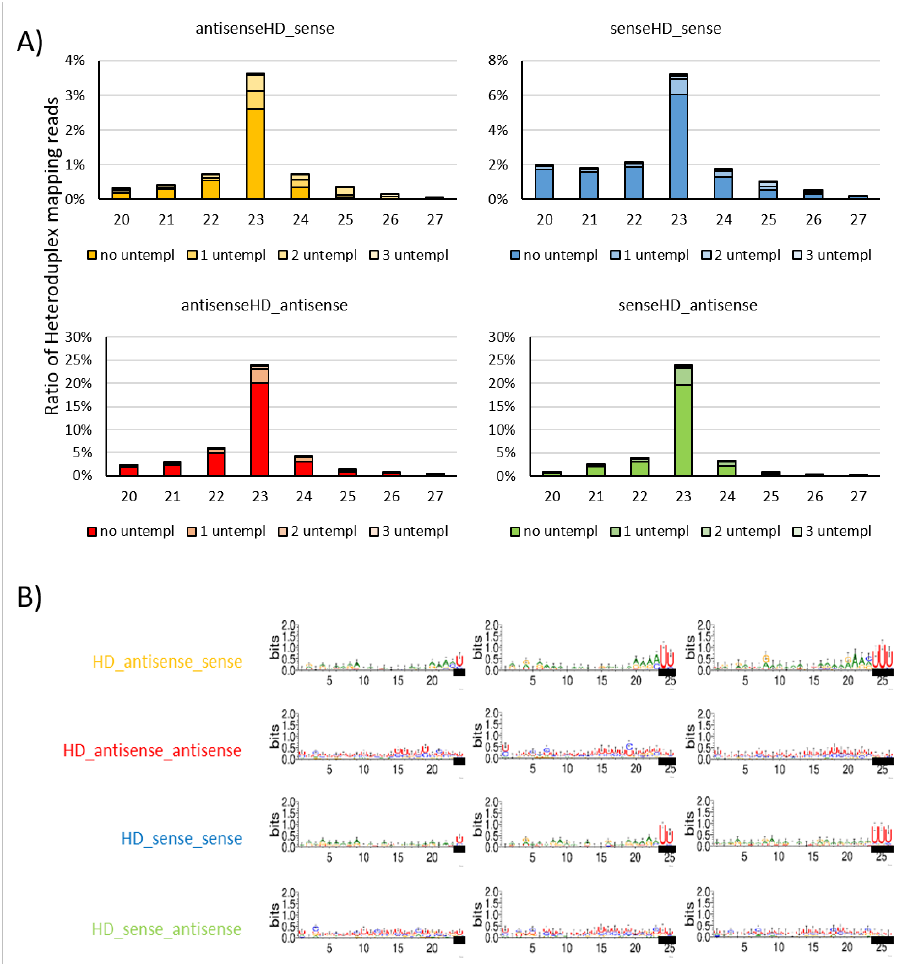
Analysis of 3′-nontemplated nucleotides in small RNAs. A) Read lenght distribution of sRNAs mapping to the individual strands. Next to percent mapping reads without mismatches, also reads are indicated with 1-3 non-templated nucleotides. B) Sequence logos of 23+1/2/3nt reads with non-templated nucleotides (black bars).

**Fig 3.**
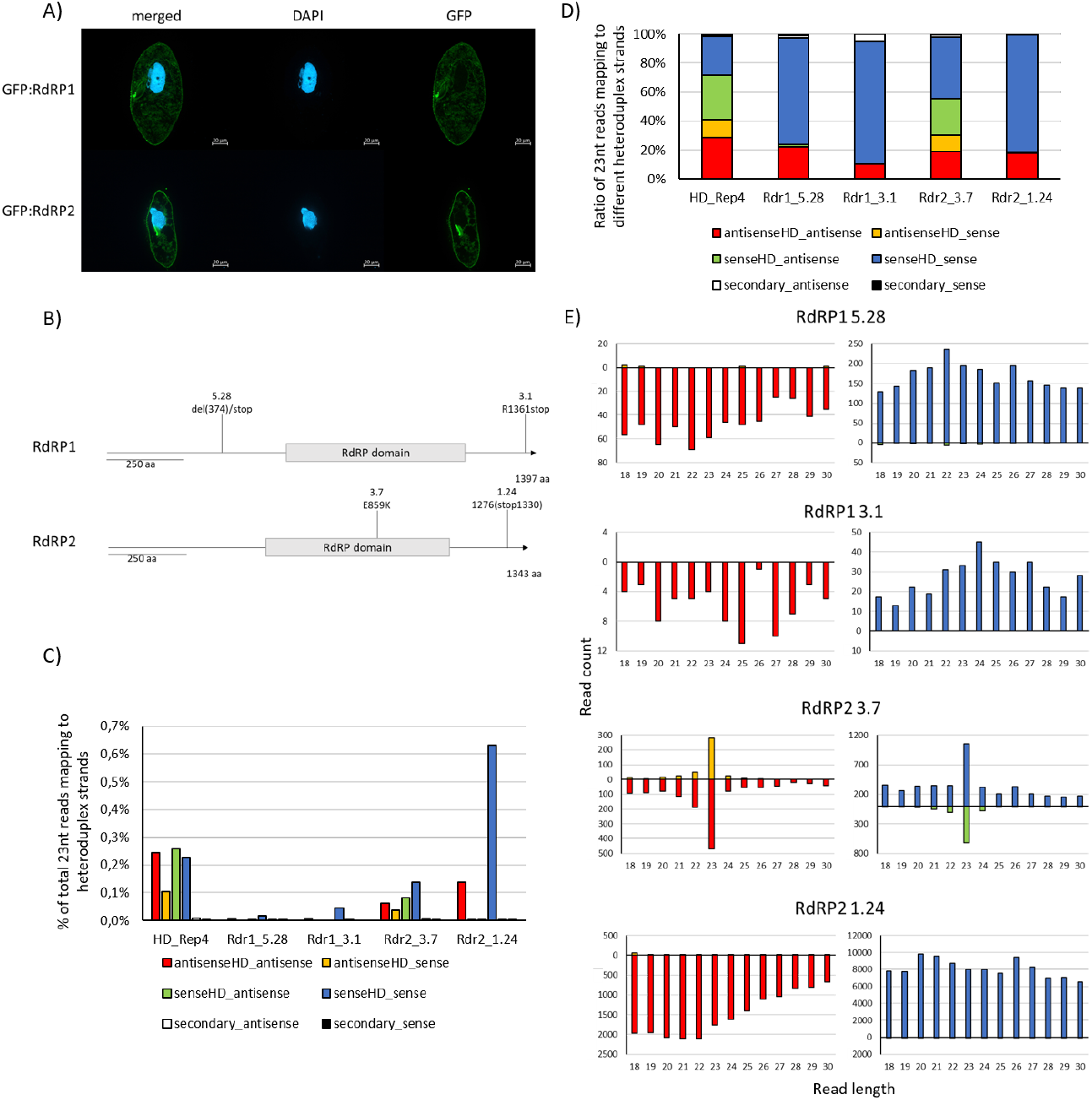
A) Immunofluorescence assay localization of GFP:RDR1 and GFP:RDR2. Fluorescence signals generated by anti-GFP antibodies are shown. DAPI is used to counterstain the macronuclei. B) Scheme of mutant cell lines used including positions annotation of mutations C) Abundance of 23nt siRNA reads mapping to the different heteroduplex strands in relation to total reads sequenced D) Ratios of 23nt siRNAs reads mapping to individual heteroduplex sequences in relation to total number of 23nt siRNA reads mapping to all heteroduplex strands E) Read length distribution of reads mapping to the different heteroduplex strands

### Exogenous dsRNA cannot be cleaved by Dicer directly

We proceeded with deeper investigations of the two RDRs. N-terminal GFP-fusion proteins of RDR1 and RDR2 co-localize in the cytosol (Fig.3A). Suppl. Fig.5 shows in addition, that we cannot observe different localizations of both RDRs when dsRNA is applied to the cells, indicating that there is no special accumulation at e.g. vacuoles when excessive dsRNA is ingested.

For functional analysis, we used mutants obtained by a forward genetic screen identifying mutants deficient for dsRNA feeding. Two RDR1 mutants have been shown to be null-alleles, RDR2 revealed hypomorphic mutants only, suggesting that this is an essential gene [37]. Fig.3B shows the positions and type of mutations in RDRs. One mutant strain of each RDR shows truncated C-termini. Structural analysis of both RDRs suggests a highly similar structure of both with the C-termini directed outward and not being associated with the catalytic core (Suppl. Fig. 6). It seems tempting to speculate that these C-termini may be necessary for complex formation.

We fed the heteroduplex particles to these strains and sequenced the resulting sRNAs. Figs. 3C-E show that no mutant line was able to produce wild type levels of 23nt siRNAs. All mutants, except RDR2-3.7, are unable to produce any RDR products and importantly also do not show 23nt siRNAs from applied dsRNA but instead only degradation products, indicating a complete lack of Dicer activity in exogenous dsRNA in RDR mutants.

We conclude that the applied dsRNA is not Dicer substrate but becomes accessible for cleavage only after being converted into new sets of dsRNA by RDRs. Both RDRs are necessary for this activity and the C-terminal region of both RDRs, which is devoid of any catalytic activity, appears to be crucial for this joint activity. In support of this, RDR2-3.7, with a mutation in the catalyic domain but with intact C-terminus, shows lower but still detectable 23nt siRNA accumulation.

If exogenous RNA was converted to dsRNA by RDRs, the next question was whether ssRNA could also trigger siRNA accumulation. Antisense 23nt siRNAs, that map to bacterial transcripts, were previously found in *Paramecium* [17]. Fig. 4 and Suppl. Fig. 7 show that both, mismatch containing sense and antisense ssRNA fed to *Paramecium*, can be efficiently converted into 23nt siRNAs, which are again accumulated predominantly from the antisense strands. This process depends on RDR1 as both mutants reveal a lack of siRNAs. In contrast to the application of dsRNA, ssRNA feeding is apparently independent of RDR2 although siRNA profiles show differences in (i) more abundant sense siRNAs and (ii) less pronounced 23nt peaks for the R2-3.7 mutant (Suppl. Fig. 7).

**Fig 4.**
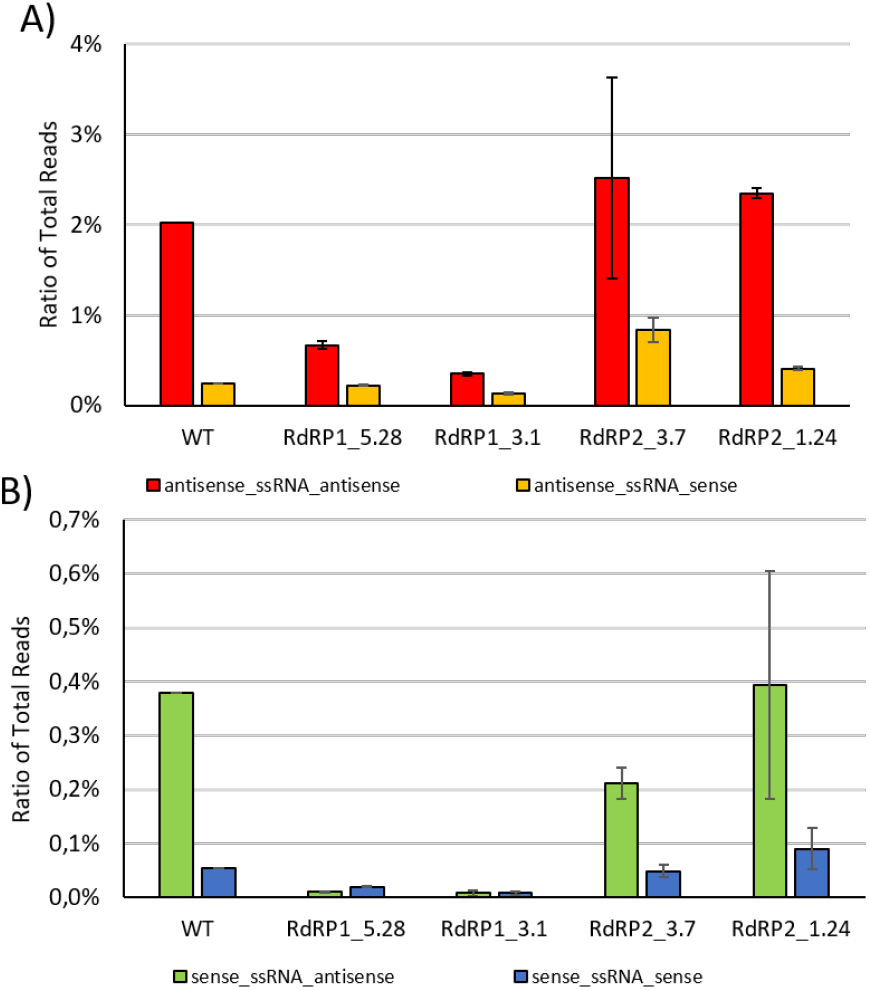
Mapping statistics of sRNA sequencing of cultures undergoing ssRNA feeding: antisense ssRNA (A) and sense ssRNA feeding (B). Only 23 nt siRNAs are quantified here.

**Fig 5.**
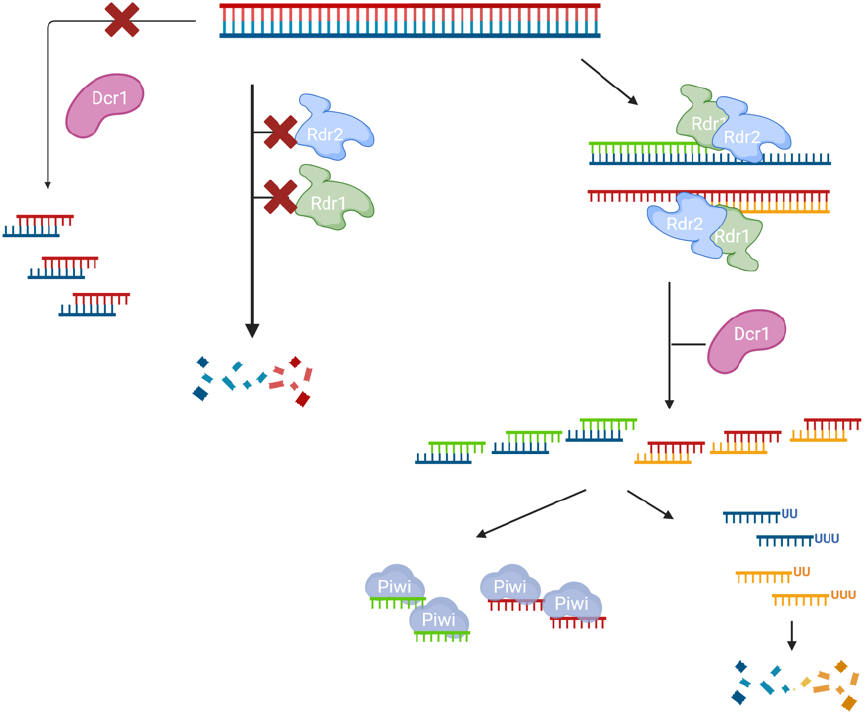
A working model for dsRNA feeding in *Paramecium*. Exogenous dsRNA cannot be cleaved by Dicer directly (left). In absence of either RDR1 or RDR2, exogenous RNA becomes degraded (middle). In WT cells, both RDRs need to interact to convert dsRNA into new hybrid-dsRNA which is Dicer substrate. It is likely that Dicer directly associates with the complex formed by both RDRs. Sense siRNA become oligouridinylated while antisense siRNAs accumulate being stabilized in Piwi proteins. (Right) Created in BioRender. Simon, M. (2025) https://BioRender.com/5f9pujd

## Discussion

Our data provide several new insights that suggest novel mechanistic roles for eukaryotic RDRs. First, RDR1 and RDR2 appear to be required to separate self-from non-self RNA and to convert this exogenous RNA into Dicer-cleavable dsRNA. It is a striking difference to the literature from other species that Dicer is not the first instance to recognize exogenous dsRNA and, in particular in relation to *C*.*elegans*, that this RDR function is also capable of converting ssRNA into siRNA [38]. In fact, our data suggest that dsRNA induced RNAi in *Paramecium* is not triggered by dsRNA itself: dsRNA delivery may simply be more efficient because the trigger is more stable than ssRNA. Our data do not indicate RNAi amplification, but rather 1:1 copying of exogenous RNA.

We hypothesize that RDR1 and RDR2 work in complexes together with Dicer; complex formation may depend on the C-terminal part of RDRs as they are necessary for activity. We see differences in RDR2 dependency between ds- and ssRNA feeding whereas RDR1 is involved in both. This means that both RDRs are not redundant but have specific roles, which is in agreement with previous reports that, for example, RDR2 is involved in transgene-induced silencing, while RDR1 is not [39].

The complexes RDRs associate in remain to be characterized. RDRCs (RNA-directed RNA polymerase complexes) in general have been described in many species, however in particular in association with endogenous RNA. In the related *Tetrahymena*, Dicer associates with RDR as shown for the production of endogenous siRNAs [40]; In yeast, RDRC is associated with heterochromatin and pericentromeric RNA and in *C*.*elegans*, an RDRC has been shown to be necessary for endogenous 26G-RNA synthesis [41]. RDRCs have yet not been reported to act on exogenous templates. It remains unknown how exogenous RNA is recognized, since both RDRs also accept endogenous RNAs and are involved in endogenous siRNA generation [42].

This new role for RDRs appears to be conserved at least in *Paramecium* spp. as phylogenetic analyses indicate orthologs also in the distant *P*.*bursaria* (Suppl. Fig. 8) agrees with earlier analyses [43]. Since other species RDR orthologs cluster mainly due to the organisms source, we cannot rule out the possibility that the role of RDRs in trigger recognition holdsother unicellular eukaryotes.

What would be the biological role of RDR mediated non-self RNA recognition and entry into the RNAi pathway? An antiviral mechanism seems logic, but it is unable to be tested since not a single virus is known to infect *Paramecium. Paramecium* in wild life is known, in particular, as a host for a great variety of endocytosymbionts, prokaryotes, and eukaryotes [44]. Recent evidence proofs an RNA-RNA interaction of *P*.*bursaria* with its facultative symbiont, the green algae chlorella, which is maintained in phagosomes: the meaning of siRNA generation from symbiont RNA may not just be in a hijacking mode in which a parasite modulates the host, but in an evolutionary conserved synchronization of metabolic integration and symbiont population control to enable a long term maintenance of the symbiotic interaction [45, 46]. RDR activity on exogenous RNA could be seen as an entry control into this mechanism, allowing the host to maintain control, in this case to prevent Dicer from uncontrolled cleavage of dsRNA or secondary structures in ssRNA.

One would also have to question the general function of 2° siRNAs in protists. In multicellular species, RDR mediated RNAi amplification means the production of large amounts of 2° siRNAs: few trigger molecules in form of 1° siRNAs cleaved from exogenous triggers are sufficient to produce many more 2° siRNAs, for instance in *C*.*elegans* by 100fold [34]. This makes sense for systemic silencing in order to disseminate an increasing number of siRNAs in multicellular organisms to achieve virus resistance in cells far away from the trigger. A single-celled species would not need such massive amplification, and thus a different mechanistic usage of RDRs seems plausible. Although 2° siRNAs exist outside the feeding region, their abundance is in single digit percentage of 1° siRNAs. This is not in the meaning of amplifying RNAi.

From the biochemical point of view, it remains unknown why two RDRs are necessary for this activity on exogenous RNA. Future studies will need to characterize the RDRC composition and biochemical activity of RDRs. Given that we see differences in dsRNA and ssRNA that feed on the dependency of RDR2 and in reference to studies in *Tetrahymena* indicating several different RDRCs associated with a single RDR [47], we expect a great complexity of different RDRCs in *Paramecium*.

## Acknowledgments

This work was supported by grants from the German Research Council (DFG) to MS (SI1379/3-1) and by the DFG Research Infrastructure NGS CC (project 407495230) as part of the Next Generation Sequencing Competence Network (project 423957469). Sample sequencing was carried out at the Competence Centre for Genomic Analysis (Kiel). M. Pirritano received a doctoral scholarship from the German Academic Scholarship Foundation (Studienstiftung des deutschen Volkes). K. Shehu received a grant from the German Academic Exchange Service (DAAD) to carry out her work (Research Grant 57552340). Yulia Yakovleva received a grant from the DAAD-Dmitrij Mendeleev Program 2021. We thank Sandra Duharcourt, Paris and Alexey Potekhin, Innsbruck for kind support with mutant strains.

