## Supplementary Data for "Exogenous dsRNA made accessible to Dicer by two eukaryotic RNA-dependent RNA polymerases"

### Suppl. Fig. 1

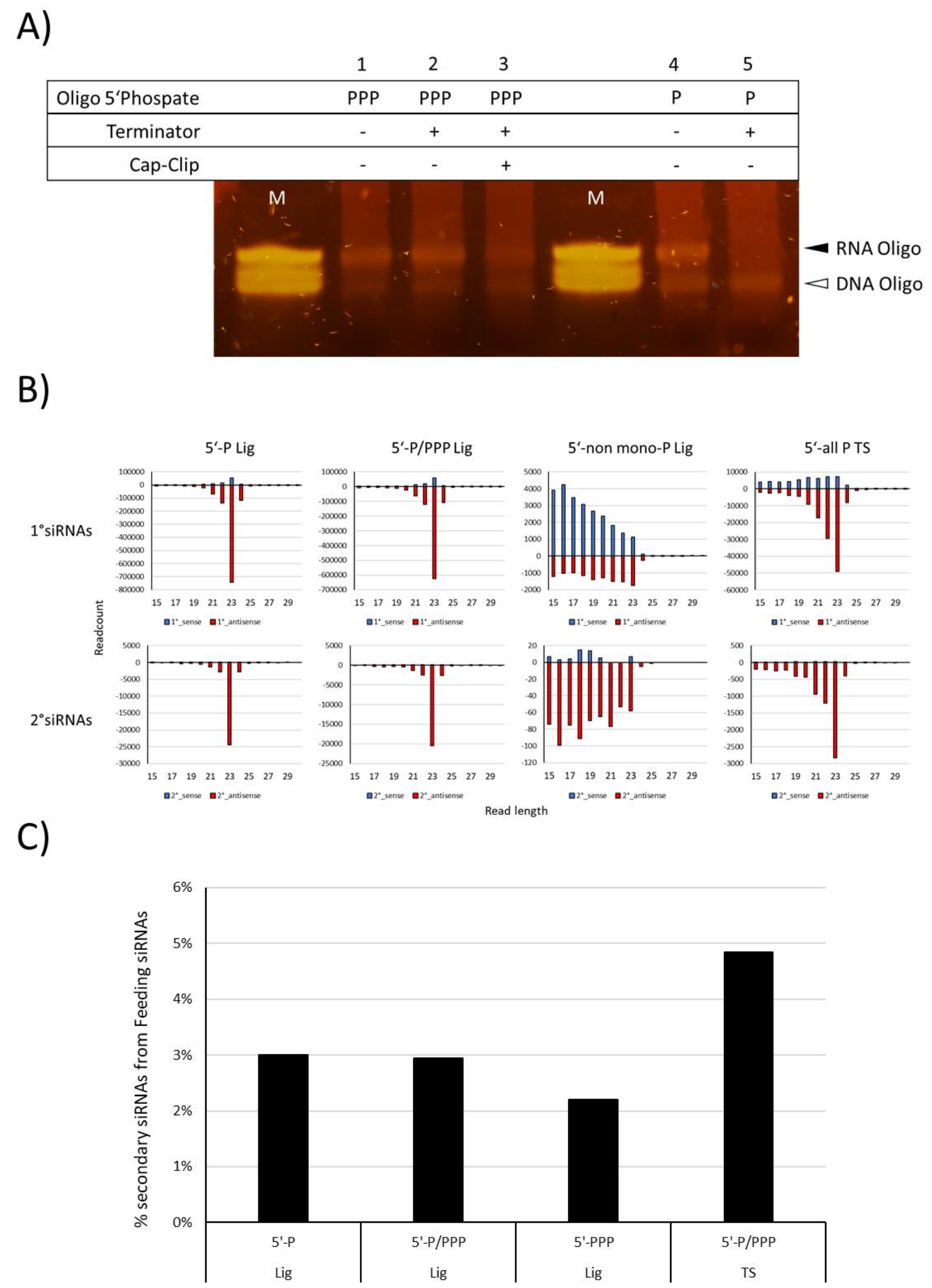


A) Analysis of control RNA oligos (5‘-tri-phosphate (left) are 5‘-mono-phosphate (right) as indicated in above line) during biochemical treatment of RNA samples. A DNA oligo resistant to all treatments was used as a loading control. Analysis on the right shows treatment with Terminator 5´-monophosphate specific exonuclease (lane 2) and subsequent treatment with CapClip pyrophosphatase to create 5´-monophosphate ends at remaining RNAs suitable for ligation (lane3). Lane 1 shows RNA sample with oligos without any treatment. Lane 4 shows the RNA sample with control oligos whithout and lane 5 with Terminator treatment. All treatments of RNA samples were carried out in duplicate, one with and one without control oligos. Parallel samples without control oligos were sequenced.

B) Read length distribution of 1° (top row) and 2° (bottom row) small RNAs derived by dsRNA feeding present in libraries of different types (Lig for ligation-based and TS for Template switch-based) enriching for different 5’ phosphorylation states (compare methods section).

C) Ratio of 2° siRNAs in relation to total feeding associated smallRNA reads within the different libraries discriminating between biochemical status of the 5’ phosphorylation. (Lig – ligation based library, TS- template switch based library)

### Suppl. Fig. 2

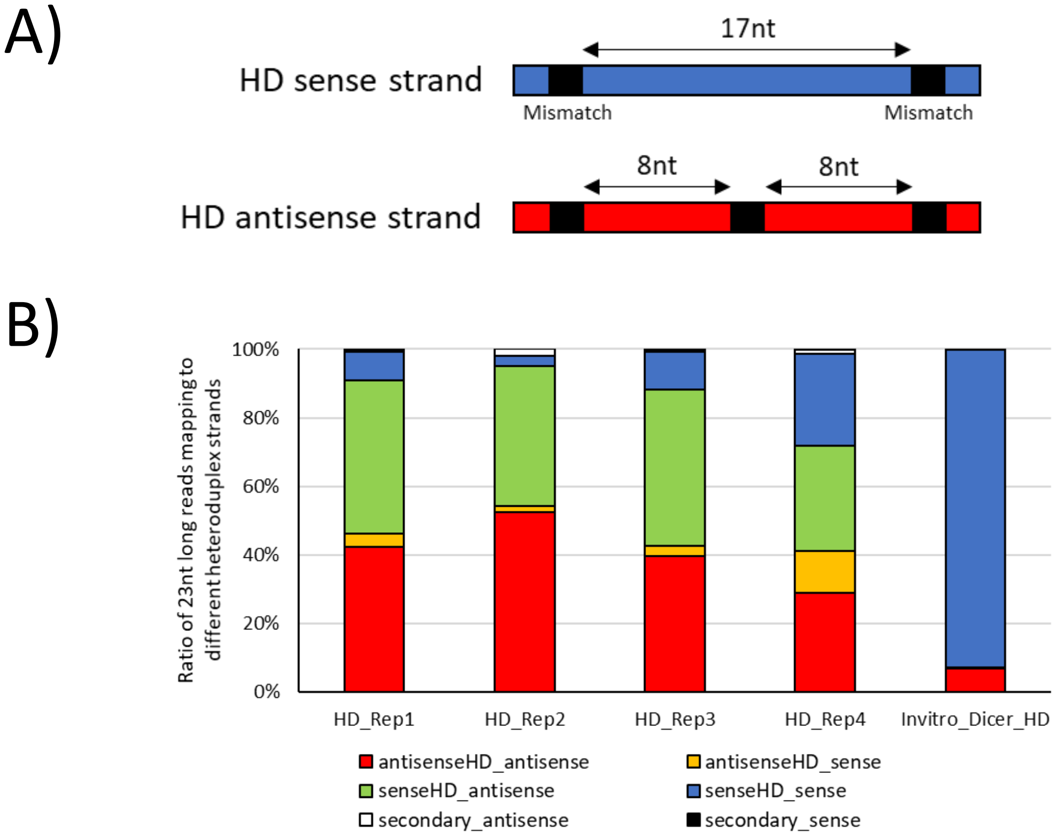


Application of heteroduplex dsRNA loaded Dextran nanoparticles

A) Schematic overview of mismatch distribution within the mismatch area of the heteroduplex dsRNA. Mismatches (black bars) and the distance between the mismatches are indicated in the sense (blue) and the antisense (red) strand of the heteroduplex.

B) Composition of 23nt siRNA reads mapping to different strands. Displayed are four biological replicates of cells fed with Heteroduplex dsRNA in addition to heteroduplex dsRNA that has been processed by an *in vitro* Dicer system instead of being applied to cells as a negative control.

### Suppl. Fig. 3

A)

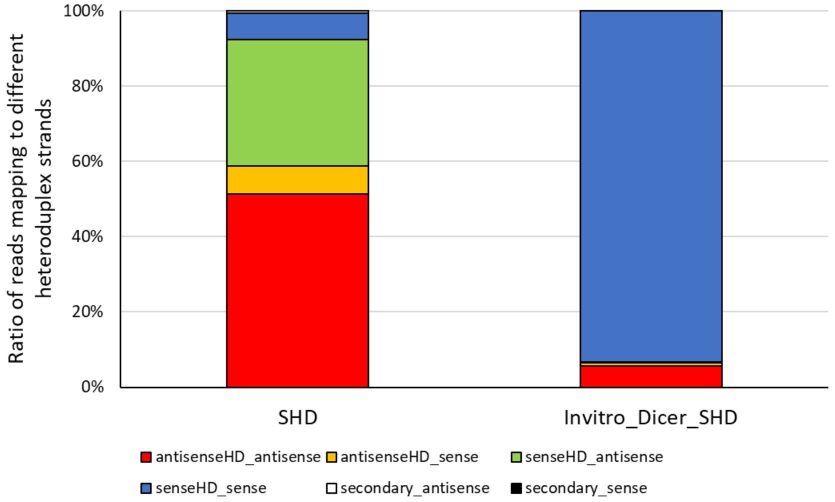


B)

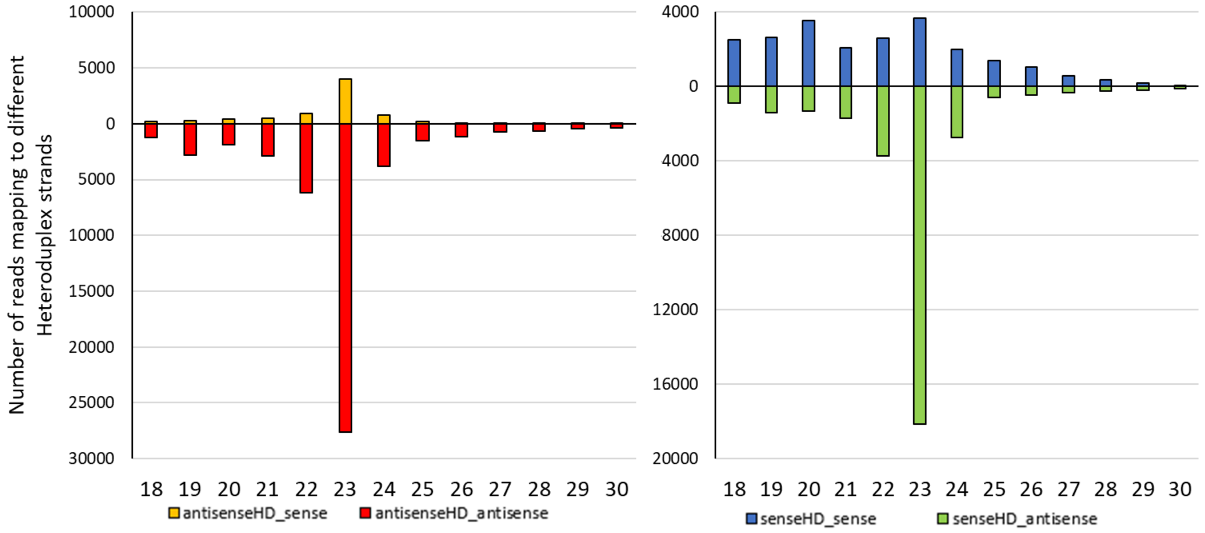


Switch heteroduplex application.

A) Ratio of 23nt siRNA reads mapping to different switch heteroduplex-associated sequences. Cells were fed with switch heteroduplex dsRNA which has the additional mismatches to dissect between both strands not in the antisense but the sense strand (blue) (Ext Fig. 2A). The same dsRNA has also been subjected for *in vitro* Dicer digestion.

B) Read length distribution of small RNA reads mapping to sense and antisense strands of the switch heteroduplex.

### Suppl. Fig. 4

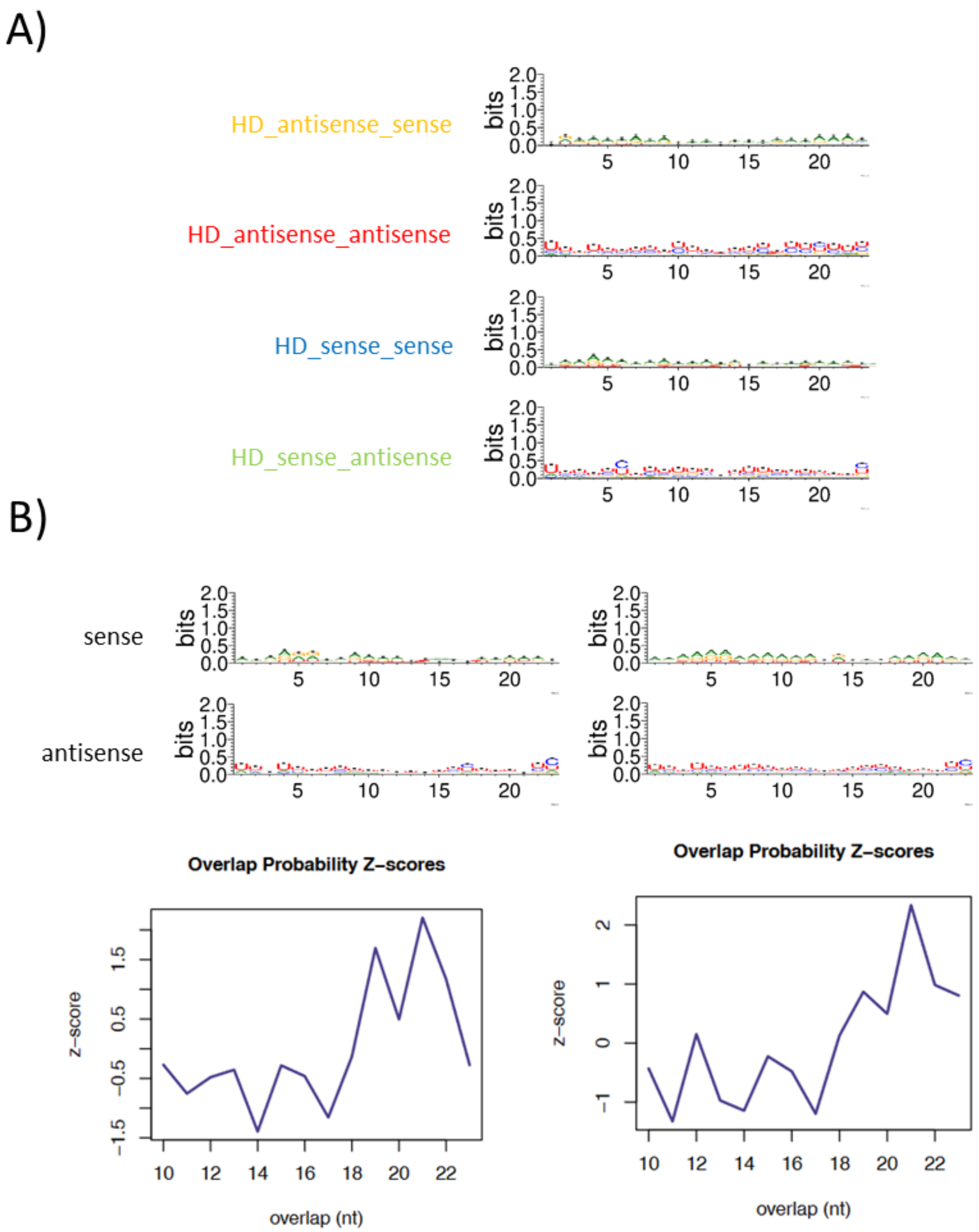


A) Sequence Logo of 23nt siRNAs produced from Heteroduplex dsRNA feeding. Displayed are the generated logos for each of the four possible Heteroduplex strands, identified using the incorporated mismatches.

B) Sequence Logo of 23nt siRNAs produced from ordinary bacteria-based dsRNA feeding. Displayed are the generated logos for sense and antisense orientated siRNAs. In addition to sequence logo generation, overlap analysis of 23nt siRNA reads was performed. Two replicates of dsRNA feeding are displayed.

### Suppl. Fig. 5

A)

RdRP1

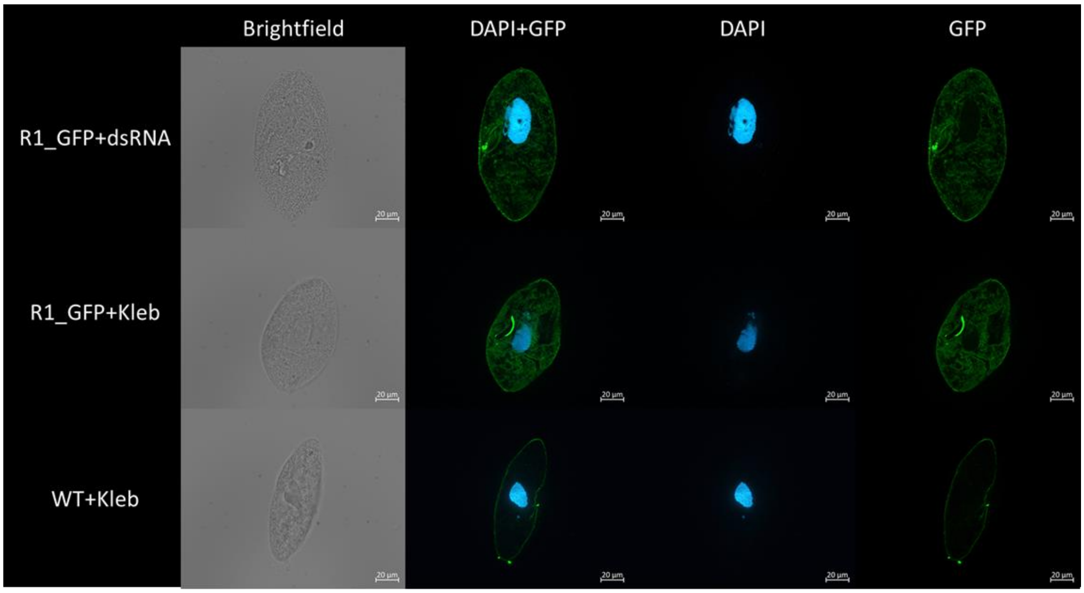


B)

RdRP2

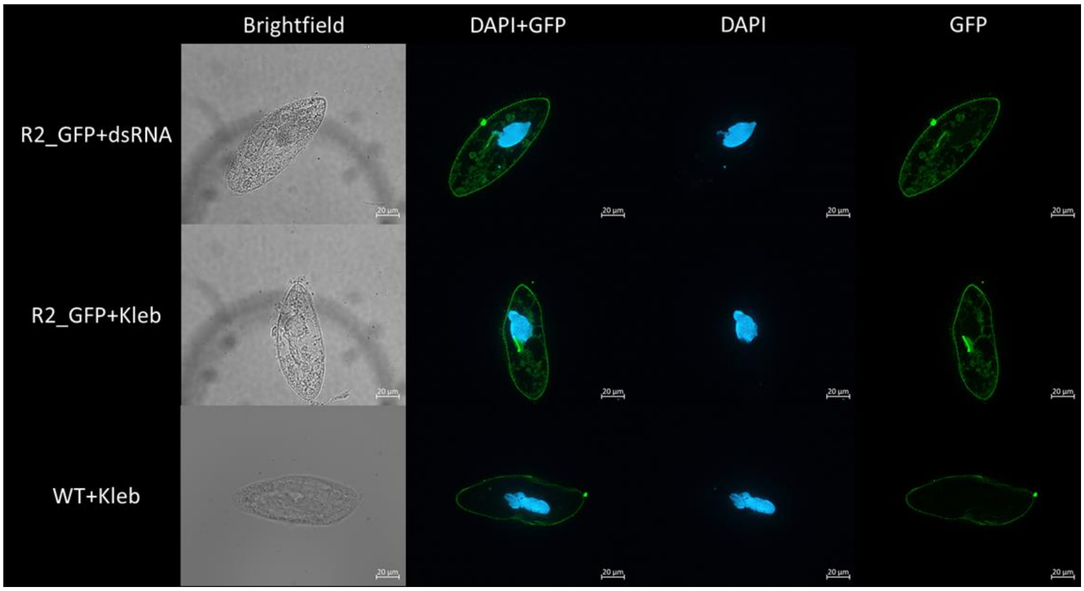


Immunofluorescence assay localization of N-terminal GFP-tagged RdRP1 and RdRP2.

Fluorescence signals were improved by anti-GFP antibodies with DAPI as counterstaining showing the macronucleus. RDR1 (A) and RDR2 (B). Brightfield images for displayed cells are provided. Localization of both RDRs was performed for both, the presence of dsRNA within food bacteria (+dsRNA) and bacteria without (+Kleb).

### Suppl. Fig. 6

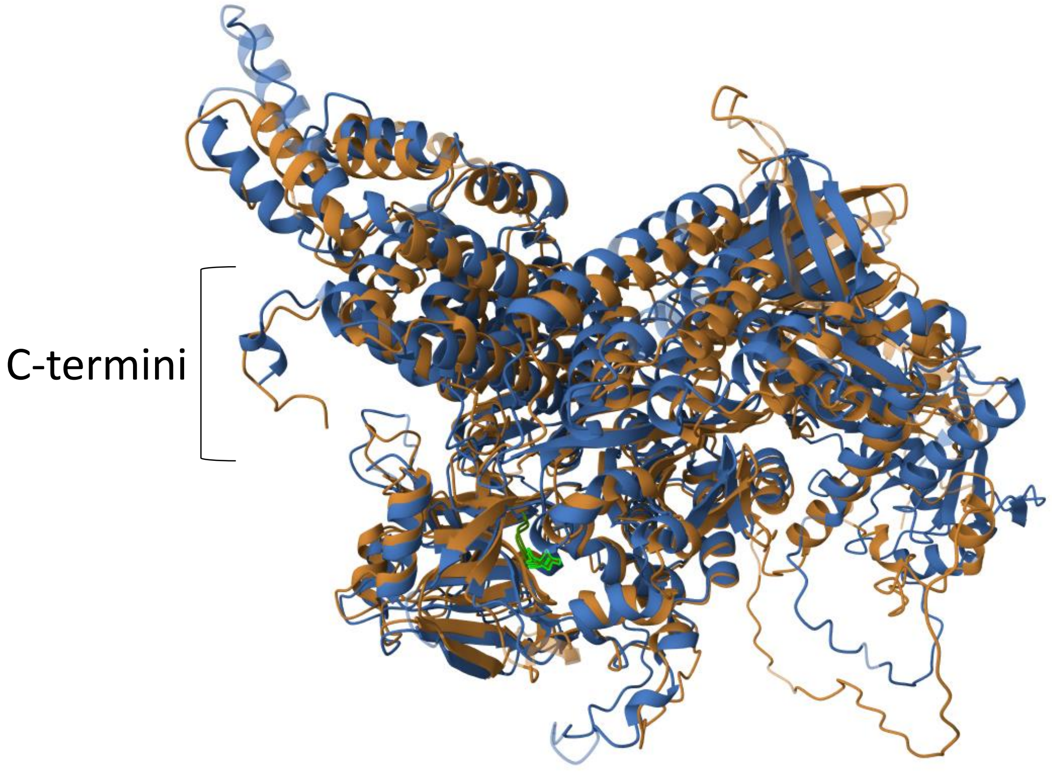


Structural alignment of Alpha Fold2 predicted structures of RDR1 (brown) and RDR2 (blue). C-terminal parts are indicated on the left and the catalytic DLDGD motif of both RDRs is indicated in green.

### Suppl. Fig 7

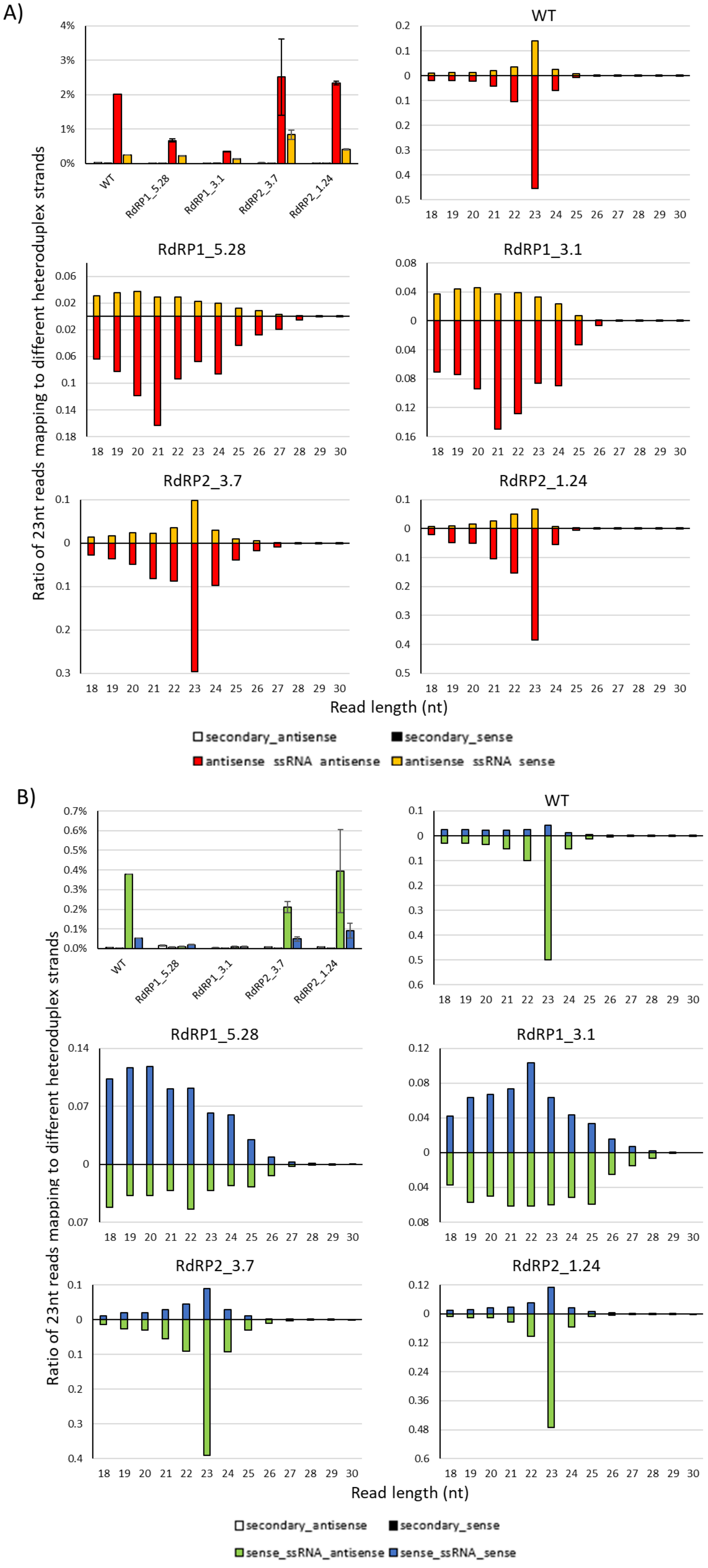


Single strand RNA feeding.

Mapping statistics and length distributions of different cell lines fed with antisense (A) or sense (B) orientated ssRNA.
Mapping statistics (first Figure in each panel) show the ratio of 23nt siRNA reads mapping to different heteroduplex related sequences in relation to total 23nt reads. Read length distributions are shown for wildtype or mutant strains, respectively.

### Suppl. Fig. 8

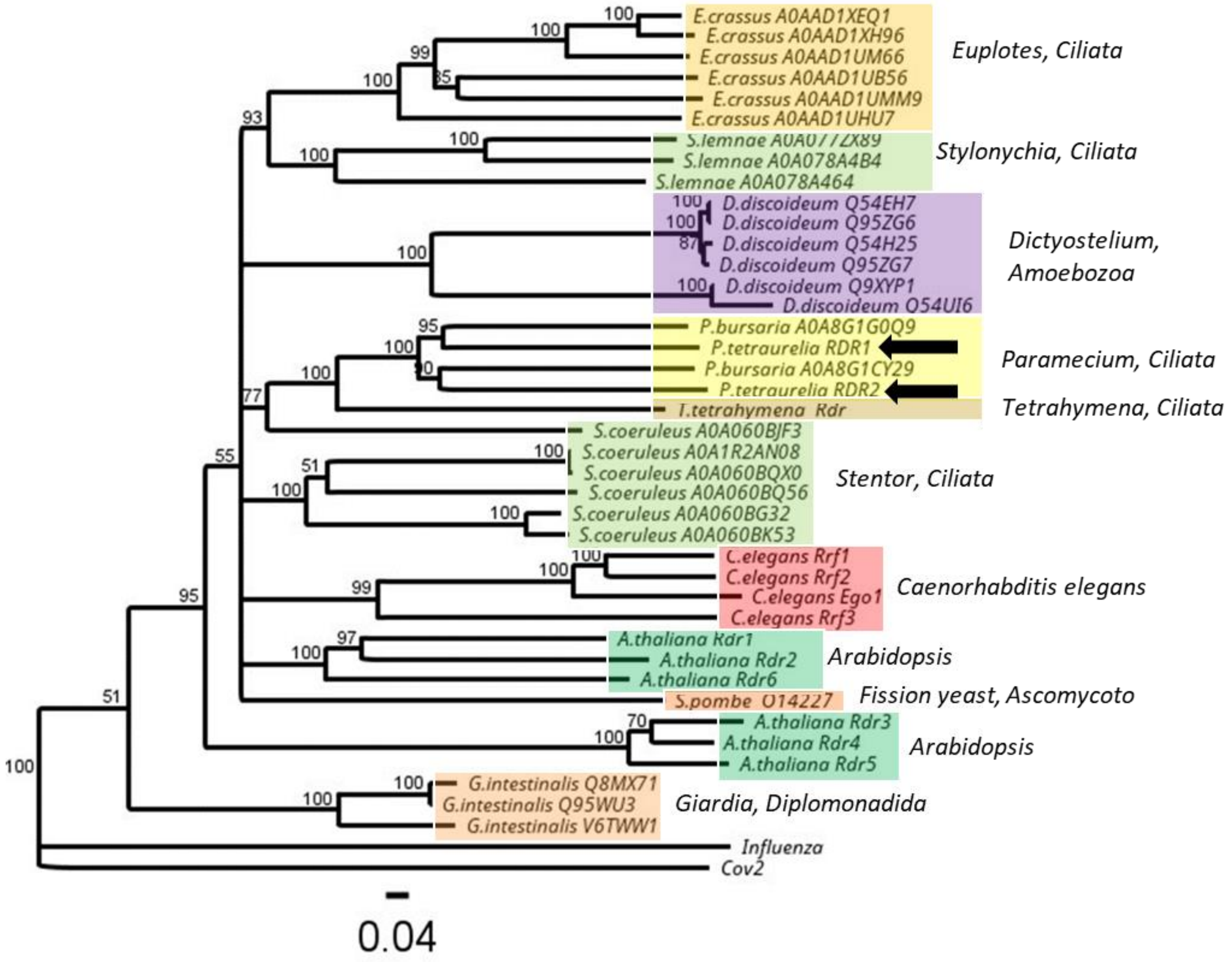


Neighbor joining consensus tree of RDR proteins rooted to the Sars-Cov-2 RDR. Supportvalues are given at the nodes.
For undesribed RDRs, UniProt Acc Nrs are indicated next to the organisms. Arrows indicate RDR1/2 from *P.tetraurelia*.
